# Age-Group Differences in the Temporal Structure of Speech and Forehead total-Hb Responses During a Phonemic Verbal Fluency Task: A Comparison of Adults in Their 40s and 70s

**DOI:** 10.64898/2026.08.22.746423

**Authors:** Kenji Nakamura

## Abstract

In populations with well-preserved cognitive function, cognitive screening total scores tend to cluster near the ceiling, making it difficult to characterize age-group differences from total scores alone. We compared the temporal structure of word production, acoustic features, and the magnitude and timing of forehead total-hemoglobin (total-Hb) responses during a phonemic verbal fluency task in adults in their 40s and 70s. A total of 254 healthy participants (115 in their 40s, 139 in their 70s) completed a 60-s phonemic verbal fluency task requiring words beginning with the Japanese syllable /ka/, administered as part of the Japanese version of the Montreal Cognitive Assessment (MoCA-J). We derived the total word count, word counts in 10-s bins, mean inter-word pause duration, speech offset time, smoothed cepstral peak prominence (CPPS), jitter, and shimmer. Area under the curve (AUC) and time-to-peak (TTP) were computed from forehead total-Hb signals recorded with a wearable single-wavelength near-infrared spectroscopy device. MoCA-J scores clustered near the ceiling in both groups, although the age-group difference was significant. The 70s group produced fewer words (14.00 ± 1.18 vs 17.09 ± 1.33) and showed longer inter-word pauses (1.60 ± 0.40 vs 0.72 ± 0.31 s). CPPS was lower, AUC was higher, and TTP was longer (32.97 ± 9.31 vs 15.63 ± 4.98 s) in the 70s group, whereas jitter did not differ. Word counts across 10-s bins showed an age group × time-bin interaction. Within each age group, participants who produced more words showed longer TTP. Age-group differences in TTP and AUC persisted after adjustment for speech offset time (proportions mediated, 6.7% and 0.5%) and in a subsample matched on speech offset time. Even when screening scores clustered at the ceiling, the temporal structure of word production and forehead total-Hb responses differed between age groups, and these two classes of measures dissociated. Because the sample was selectively recruited and single-wavelength total-Hb signals do not index localized neural activity, the findings are descriptive and motivate longitudinal, multi-axis characterization of speech in aging.

## 1 Introduction

Age-related changes in cognitive function vary substantially among individuals; even within the same age band, some people maintain cognitive function relatively well whereas others decline (Cabeza et al., 2018). Longitudinal multimodal biological data further show that different biological systems can change at different rates and in different directions within the same individual, which limits the usefulness of characterizing aging with any single measure (Ahadi et al., 2020). Cognitive screening instruments such as the Montreal Cognitive Assessment (MoCA) are clinically useful, but their restricted score range means that distributions cluster near the ceiling in populations with well-preserved cognition, so that differences in how a task is performed may not be represented by the total score. There is therefore value in evaluating the temporal structure of task performance and concurrent physiological responses alongside total scores.

The phonemic verbal fluency task requires participants to generate as many words as possible beginning with a specified sound within a fixed period, and it engages lexical access, retrieval, selection, inhibition, and speech production. Phonemic and semantic fluency follow different lifespan trajectories: decline in later life is more pronounced for semantic fluency, whereas phonemic fluency is relatively preserved (Kavé and Knafo-Noam, 2015). Phonemic fluency is nonetheless affected by aging, and normative data from a large community cohort indicate a decline of approximately one word per decade (Vaughan et al., 2016). Although the total number of words produced in one minute is the conventional outcome, dividing the minute into shorter intervals shows that the within-task gradient of word production differs by cognitive status and carries information distinct from the total count (Holtzer et al., 2020). That study, however, compared participants with and without mild cognitive impairment and did not examine age differences among healthy adults. Lexical search during verbal fluency has also been modeled as a dynamic process in which local exploitation of activated regions alternates with exploration of new regions (Hills et al., 2012); because this model was developed primarily for semantic fluency, it informs rather than defines the analysis of phonemic tasks. Evaluating word counts in 10-s bins and inter-word pause durations, in addition to the total count, may therefore provide a more detailed characterization of age-group differences in the temporal structure of word production.

Near-infrared spectroscopy (NIRS) enables noninvasive measurement of hemodynamic signals during task performance. In verbal fluency tasks using multi-wavelength NIRS, the integral of the oxygenated hemoglobin response and the temporal centroid have been used to characterize response magnitude and timing separately (Takizawa et al., 2014). BOLD-fMRI studies report prolongation of the time-to-peak of the hemodynamic response function with healthy aging (West et al., 2019), and hemodynamic latency may be associated with cognitive ability (Anderson et al., 2020). These measures are not identical to the total-Hb signal obtained from the single-wavelength device used here. We therefore computed the area under the curve (AUC) and the time-to-peak (TTP) of the forehead total-Hb signal to characterize the magnitude and the timing of the device output separately.

Verbal fluency tasks also yield information from the speech signal itself. Several acoustic features vary with age and sex (Taylor et al., 2020), and smoothed cepstral peak prominence (CPPS) has been systematically evaluated as an index of voice quality and periodicity (Yanushevskaya and Kenny, 2025). CPPS levels differ systematically by sex, age, speech task (sustained vowel versus connected speech), and analysis software, so comparisons require standardized task conditions (Murton et al., 2020; Buckley et al., 2023). In middle-aged and older adults, voice features such as jitter have been associated with long-term cognitive change (Mahon and Lachman, 2022), and artificial-intelligence-based voice biomarkers detect cognitive impairment among community-dwelling adults in Japan (Kiyoshige et al., 2025). Spontaneous speech has also been used to identify declines in physical function, including gait, strength, balance, and body composition (Da Cunha et al., 2026). Together these findings suggest that speech carries both peripheral (physical) contributions, such as phonatory and respiratory function, and central (cognitive-linguistic) contributions, such as lexical retrieval and executive control; how far these can be separated using a single voice metric remains unresolved.

Accordingly, we administered a 60-s phonemic verbal fluency task using the Japanese syllable /ka/ to adults in their 40s and 70s and measured voice quality, task performance, speech temporal structure, and forehead total-Hb responses simultaneously from a single speech sample. Our aims were to compare these measures between age groups, to test whether the association between TTP and total word count differed by age group, and to determine whether any age-group differences in total-Hb measures could be explained by differences in how long speech continued during the task.

## 2 Materials and Methods

### 2.1 Participants and Recruitment

A total of 254 participants were analyzed: 115 adults in their 40s (40 men, 75 women; mean age 43.16 ± 2.28 years) and 139 adults in their 70s (45 men, 94 women; mean age 73.80 ± 2.59 years). The 40s group comprised staff employed at care houses (keihi rojin home, Type C low-cost residential homes for older adults in Japan), and the 70s group comprised residents of care houses of the same type. Care houses are intended for older adults who can manage personal activities independently but have concerns about living alone; people who require continuous care, or who cannot sustain communal living because of dementia or comparable conditions, are not eligible for admission. The 70s group therefore consisted of older adults who were independent in activities of daily living and had no evident cognitive impairment.

Participants were recruited through notices posted within the facilities. The notices and the accompanying explanations stated that eligible participants had to be healthy individuals with none of the following: a history of central nervous system or psychiatric disease; a voice disorder; hearing or visual impairment severe enough to interfere with task performance; a non-native Japanese language background; a chronic condition requiring regular medical visits or treatment (for example hypertension, diabetes, or cardiovascular disease); or ongoing use of vasoactive medication. Eligibility was confirmed individually for each applicant before the study was explained, and the 254 individuals who provided written informed consent were enrolled. Eligibility was thus determined during recruitment and application: no applicant was found ineligible at the eligibility-confirmation stage, and no participant was excluded after consent. The recruitment and analysis flow is shown in Supplementary Figure S2.

All measurements were performed by a single researcher using a standardized procedure. Both groups had a pre-task rest period and were assessed in a quiet room using the same protocol. The overall study period was November 2024 to November 2025; recruitment and data collection were completed between November 2024 and March 2025. Cognitive function was assessed with the Japanese version of the MoCA (MoCA-J), the Japanese adaptation of the original MoCA (Nasreddine et al., 2005), whose reliability and validity have been established (Fujiwara et al., 2010). Years of education were not collected; consequently the one-point adjustment specified in the MoCA scoring rules for individuals with 12 or fewer years of education was not applied, and education could not be entered as a covariate.

### 2.2 Phonemic Verbal Fluency Task and Speech Analysis

We analyzed the phonemic verbal fluency item administered as part of the MoCA-J. Participants were asked to produce as many words as possible beginning with the Japanese syllable /ka/ within 60 s while speech and forehead NIRS signals were recorded simultaneously. The number of valid words was counted for each 10-s bin (0-10, 10-20, 20-30, 30-40, 40-50, and 50-60 s), and the total word count was the sum across the six bins; identical validity criteria were applied to both. Mean inter-word pause duration was defined, for adjacent valid words, as the silent interval from the offset of the preceding word to the onset of the following word, averaged within each participant; the latency from task onset to the first word and the silence after the final word were excluded. To quantify how long speech continued during the task, speech offset time was defined as the end of the last 10-s bin in which at least one word was produced. Because this definition rests on 10-s resolution, it is an upper-bound approximation of the actual offset of the final word.

Speech was recorded in a quiet room with a Fairy Devices microphone placed approximately 2 m from the participant across a desk, and stored as monaural WAV files at 44.1 kHz and 16-bit PCM. Acoustic analyses were performed with Parselmouth 0.4.3 (Praat 6.3 series; Praat API for Python) on voiced segments. Periodic points were extracted with “To PointProcess (periodic, cc)” using a pitch floor of 75.0 Hz, a pitch ceiling of 500.0 Hz, a period floor of 0.0001 s, and a period ceiling of 0.0200 s. CPPS was computed from the power cepstrogram with a pitch floor of 60 Hz, a time step of 0.002 s, a maximum frequency of 5000 Hz, and pre-emphasis from 50 Hz. The Get CPPS settings were: subtract trend = no; time averaging window = 0.01 s; quefrency averaging window = 0.001 s; peak search range = 60-330 Hz; tolerance = 0.05; interpolation = parabolic; tilt line trend type = straight; regression method = robust (Theil’s robust fit). Jitter (local) used a period floor of 0.0001 s, a period ceiling of 0.0200 s, and a maximum period factor of 1.3; shimmer (local) used the same period settings with a maximum period factor of 1.3 and a maximum amplitude factor of 1.6. Both were multiplied by 100 and expressed as percentages. CPPS was treated as the principal acoustic measure, whereas jitter and shimmer were treated as exploratory because they are more sensitive to voiced-segment extraction in connected speech.

### 2.3 Forehead NIRS Measurement

Forehead NIRS was recorded with the wearable HOT-2000 device (NeU Corporation). The HOT-2000 has two channels (left and right), uses a single wavelength (810 nm), and samples at 10 Hz; it cannot separate oxygenated from deoxygenated hemoglobin. We analyzed the relative change in forehead total-Hb reported by the device (Takahashi et al., 2022). For each participant, the mean value over the 10 s preceding task onset (-10 to 0 s) was subtracted as baseline, and a fourth-order zero-phase Butterworth low-pass filter (cutoff 0.1 Hz; forward-backward filtering) was applied. TTP was defined as the time at which the baseline-corrected, filtered total-Hb signal reached its maximum within 0-60 s after task onset; no participant peaked within 0-5 s or 50-60 s. Sensitivity analyses performed separately for the left and right channels showed the same direction of the age-group difference, with longer TTP in the 70s group. AUC was obtained by integrating the baseline-corrected total-Hb change from task onset to 60 s and is expressed in a.u.·s. Because the HOT-2000 output is a relative change, age-group differences in AUC were interpreted as differences in task-related device output rather than in absolute cerebral blood flow. The centroid value used in NIRS studies of verbal fluency is not mathematically identical to the TTP used here (Takizawa et al., 2014). During measurement no abrupt head movement or turning was observed, and the concurrently displayed pulse indicator showed no abrupt increases; however, quantitative artifact rejection using accelerometry or pulse signals and physiological regression correction were not performed. The signals were therefore interpreted specifically as task-related forehead total-Hb responses rather than as localized cortical neural activity.

### 2.4 Statistical Analysis

Continuous variables are presented as mean ± standard deviation. The primary outcomes were total word count and TTP; CPPS, jitter, shimmer, and AUC were secondary or exploratory outcomes. Groups were compared with Welch’s t test, and Hedges’ g with 95% confidence intervals was calculated as the effect size. Because acoustic and hemodynamic measures show systematic sex differences, multiple regression analyses with age group and sex as explanatory variables were performed for all outcomes. Education could not be adjusted for because years of education were not collected. Because MoCA-J scores clustered near the ceiling, the Mann-Whitney U test was used as a distribution-sensitive sensitivity analysis; because the verbal fluency item analyzed here contributes to the MoCA-J total, the same test was also applied to a modified MoCA-J score excluding that item. The Benjamini-Hochberg procedure was applied to between-group comparisons of the secondary and exploratory outcomes (CPPS, jitter, shimmer, and AUC) to control the false discovery rate (FDR). All tests were two-sided with a significance level of 5%.

Word counts in each 10-s bin were treated as count data and analyzed with generalized estimating equations (GEE) accounting for six repeated measurements per participant, using a Poisson family, a log link, an exchangeable working correlation structure, and robust sandwich standard errors. The model included age group, time bin, and their interaction. The Pearson dispersion statistic was approximately 0.30, indicating underdispersion; robust standard errors were therefore used. Associations between NIRS measures and verbal fluency performance were quantified with Pearson product-moment correlations, followed by analyses adjusting for age group and sex. Because the direction of the TTP-word count correlation differed between the pooled and age-stratified analyses, an additional regression analysis was performed with total word count as the dependent variable and age group, TTP, their interaction, and sex as explanatory variables.

Three further analyses tested whether age-group differences in forehead total-Hb responses could be explained by between-group differences in how long speech continued. First, age-group differences in TTP and AUC were re-estimated with speech offset time and sex as covariates. Second, mediation analyses were conducted with age group as the independent variable, speech offset time as the mediator, and TTP or AUC as the dependent variable; 95% confidence intervals for indirect effects were obtained by bootstrap resampling with 5,000 iterations. Third, a subsample analysis was restricted to participants who continued speaking through the final 10-s bin, so that speech offset time was identical in the two groups. Analyses were performed in Python 3.13.5 with SciPy 1.17.0 and statsmodels 0.14.6.

## 3 Results

### 3.1 Between-Group Comparison of Primary and Secondary Measures

Sex distribution did not differ between age groups (men: 34.8% in the 40s group, 32.4% in the 70s group; χ2 = 0.074, p = 0.786). The MoCA-J total score was 29.96 ± 0.20 in the 40s group and 29.78 ± 0.51 in the 70s group, and the age-group difference was significant by both Welch’s t test (t = 3.66, df = 188.85, p < 0.001) and the Mann-Whitney U test (U = 9040, p = 0.001). The median [interquartile range] was nevertheless 30 [30-30] in both groups, and 95.7% of the 40s group and 82.7% of the 70s group obtained a perfect score, indicating a strong ceiling concentration. Observed score ranges were 29-30 in the 40s group (110 participants scored 30 and 5 scored 29) and 28-30 in the 70s group (115 scored 30, 18 scored 29, and 6 scored 28). In the sensitivity analysis using the modified MoCA-J score excluding the verbal fluency item, no age-group difference was observed at the two-sided 5% level.

The total number of words produced was 17.09 ± 1.33 in the 40s group and 14.00 ± 1.18 in the 70s group, a difference of 3.09 fewer words in the 70s group (95% CI -3.40 to -2.77; Hedges’ g = -2.46; p < 0.001). Mean inter-word pause duration was 0.72 ± 0.31 s and 1.60 ± 0.40 s, respectively, 0.88 s longer in the 70s group (95% CI 0.80 to 0.97; Hedges’ g = 2.44; p < 0.001).

CPPS was 15.18 ± 1.25 dB in the 40s group and 12.91 ± 1.19 dB in the 70s group, lower in the 70s group (FDR-adjusted q < 0.001). Jitter was 2.18 ± 0.18% and 2.23 ± 0.32%, with no significant age-group difference (q = 0.117). Shimmer was 21.37 ± 1.90% and 20.43 ± 1.69%, lower in the 70s group (q < 0.001). Clear sex differences were present in the acoustic measures: in sex-adjusted regression, CPPS was 1.59 dB higher in men (p < 0.001) and jitter was 0.27% lower in men (p < 0.001). Sex-stratified CPPS values were 14.57 ± 0.97 dB in women and 16.32 ± 0.86 dB in men in the 40s group, and 12.44 ± 0.85 dB in women and 13.89 ± 1.22 dB in men in the 70s group. Age-group differences persisted after adjustment for sex (CPPS, β = -2.23 dB, 95% CI -2.47 to -1.99, p < 0.001; shimmer, β = -0.92%, 95% CI -1.37 to -0.48, p < 0.001), whereas jitter again showed no significant difference (β = 0.04%, 95% CI -0.01 to 0.10, p = 0.137).

TTP was 15.63 ± 4.98 s in the 40s group and 32.97 ± 9.31 s in the 70s group, 17.34 s longer in the 70s group (95% CI 15.54 to 19.15; Hedges’ g = 2.26; p < 0.001). AUC was 0.776 ± 0.188 a.u.·s and 1.059 ± 0.203 a.u.·s, 0.283 a.u.·s higher in the 70s group (95% CI 0.235 to 0.332; Hedges’ g = 1.44; FDR-adjusted q < 0.001). Age-group differences persisted in sex-adjusted regression for total word count (β = -3.09 words, 95% CI -3.40 to -2.78, p < 0.001), TTP (β = 17.32 s, 95% CI 15.42 to 19.22, p < 0.001), and AUC (β = 0.286 a.u.·s, 95% CI 0.240 to 0.333, p < 0.001). AUC additionally showed a main effect of sex, being 0.129 a.u.·s higher in men (p < 0.001). The main results are summarized in Table 1.

**Table 1.** Participant characteristics and outcome measures by age group.

| Variable | 40s (n = 115) | 70s (n = 139) | Mean difference (95% CI) | Hedges' g (95% CI) | p / FDR q |
| --- | --- | --- | --- | --- | --- |
| Participants, n | 115 | 139 |  |  |  |
| Male sex, n (%) | 40 (34.8) | 45 (32.4) |  |  | 0.786 |
| Age, years | 43.16 ± 2.28 | 73.80 ± 2.59 |  |  |  |
| MoCA-J score | 29.96 ± 0.20 | 29.78 ± 0.51 | -0.17 (-0.27, -0.08) | -0.43 (-0.68, -0.18) | 0.001 |
| Total words, n | 17.09 ± 1.33 | 14.00 ± 1.18 | -3.09 (-3.40, -2.77) | -2.46 (-2.79, -2.14) | < 0.001 |
| Mean inter-word pause duration, s | 0.72 ± 0.31 | 1.60 ± 0.40 | 0.88 (0.80, 0.97) | 2.44 (2.11, 2.76) | < 0.001 |
| Speech offset time, s | 53.22 ± 5.22 | 57.27 ± 4.63 | 4.05 (2.82, 5.28) | 0.82 (0.56, 1.07) | < 0.001 |
| CPPS, dB | 15.18 ± 1.25 | 12.91 ± 1.19 | -2.27 (-2.57, -1.96) | -1.86 (-2.15, -1.56) | < 0.001 |
| Jitter, % | 2.18 ± 0.18 | 2.23 ± 0.32 | 0.05 (-0.01, 0.11) | 0.19 (-0.06, 0.44) | 0.117 |
| Shimmer, % | 21.37 ± 1.90 | 20.43 ± 1.69 | -0.93 (-1.38, -0.49) | -0.52 (-0.77, -0.27) | < 0.001 |
| Time-to-peak (TTP), s | 15.63 ± 4.98 | 32.97 ± 9.31 | 17.34 (15.54, 19.15) | 2.26 (1.94, 2.57) | < 0.001 |
| AUC, a.u.·s | 0.776 ± 0.188 | 1.059 ± 0.203 | 0.283 (0.235, 0.332) | 1.44 (1.16, 1.72) | < 0.001 |

Values are mean ± SD unless otherwise noted. Mean differences and Hedges' g are oriented as 70s minus 40s. P values are from Welch two-sample t tests for continuous outcomes, except MoCA-J (Mann-Whitney U test) and sex (chi-square test). The primary outcomes were total words and TTP. Benjamini-Hochberg FDR correction was applied to the secondary and exploratory outcomes (CPPS, jitter, shimmer, and AUC). AUC is expressed in arbitrary units × seconds (a.u.·s) and is not an absolute cerebral blood-flow measure. Sex-adjusted between-group estimates for all outcomes are reported in the Results. CPPS, smoothed cepstral peak prominence; AUC, area under the curve; TTP, time-to-peak; CI, confidence interval.

### 3.2 Temporal Course of Word Production

Word production over the 1-min task was analyzed in 10-s bins. Mean word counts in the 40s group were 3.55 ± 0.50, 3.50 ± 0.50, 3.48 ± 0.52, 3.34 ± 0.51, 2.52 ± 0.93, and 0.70 ± 1.04. The corresponding values in the 70s group were 3.40 ± 0.49, 2.31 ± 0.46, 2.45 ± 0.50, 2.24 ± 0.45, 2.12 ± 0.56, and 1.48 ± 1.20. GEE analysis showed a significant age group × time-bin interaction in addition to a main effect of time bin (χ2(5) = 274.9, p < 0.001). The 40s group produced relatively more words during the first half of the task and then declined steeply, whereas the 70s group settled at a lower rate from 10 s onward and sustained it. The 1-min production profile therefore differed by age group (Figure 1).

**Figure 1.**
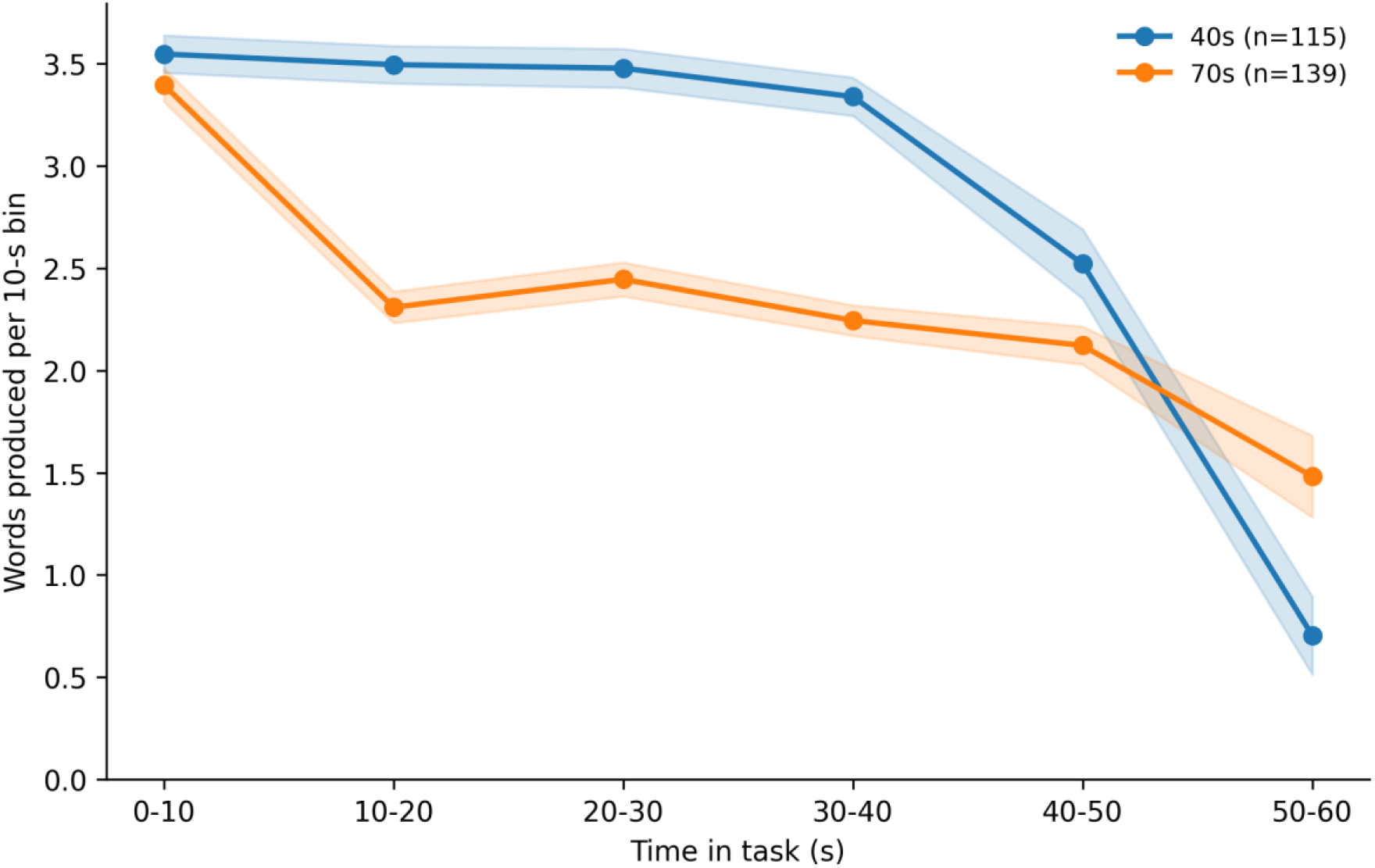
Time course of word production across the 60-s phonemic verbal fluency task. Mean number of valid words produced in each consecutive 10-s bin (0-10, 10-20, 20-30, 30-40, 40-50, and 50-60 s) for the 40s group (n = 115) and the 70s group (n = 139). The horizontal axis gives the 10-s bin and the vertical axis the number of valid words produced within that bin; bin-specific counts sum to the total word count reported in Table 1. Points denote group means and shaded bands denote 95% confidence intervals of the mean. Counts were analyzed as count data using generalized estimating equations with a Poisson family, a log link, an exchangeable working correlation structure, and robust sandwich standard errors; the age group × time-bin interaction was significant (χ2(5) = 274.9, p < 0.001). The 40s group produced more words early in the task and then declined steeply, falling to 0.70 ± 1.04 words in the final bin, whereas the 70s group settled at a lower rate from 10 s onward and sustained production through to the end of the task (1.48 ± 1.20 words in the final bin).

### 3.3 Forehead total-Hb Responses and Their Association with Word Production

The group-averaged, baseline-corrected total-Hb time course is shown in Figure 2. The 40s group reached its maximum relatively early after task onset, whereas the peak occurred later in the 70s group; mean TTP was 15.63 s and 32.97 s, respectively. Figure 2 shows group-mean waveforms with 95% confidence intervals across the 10-s pre-task baseline, the 0-60 s task phase, and the 60-80 s recovery period.

**Figure 2.**
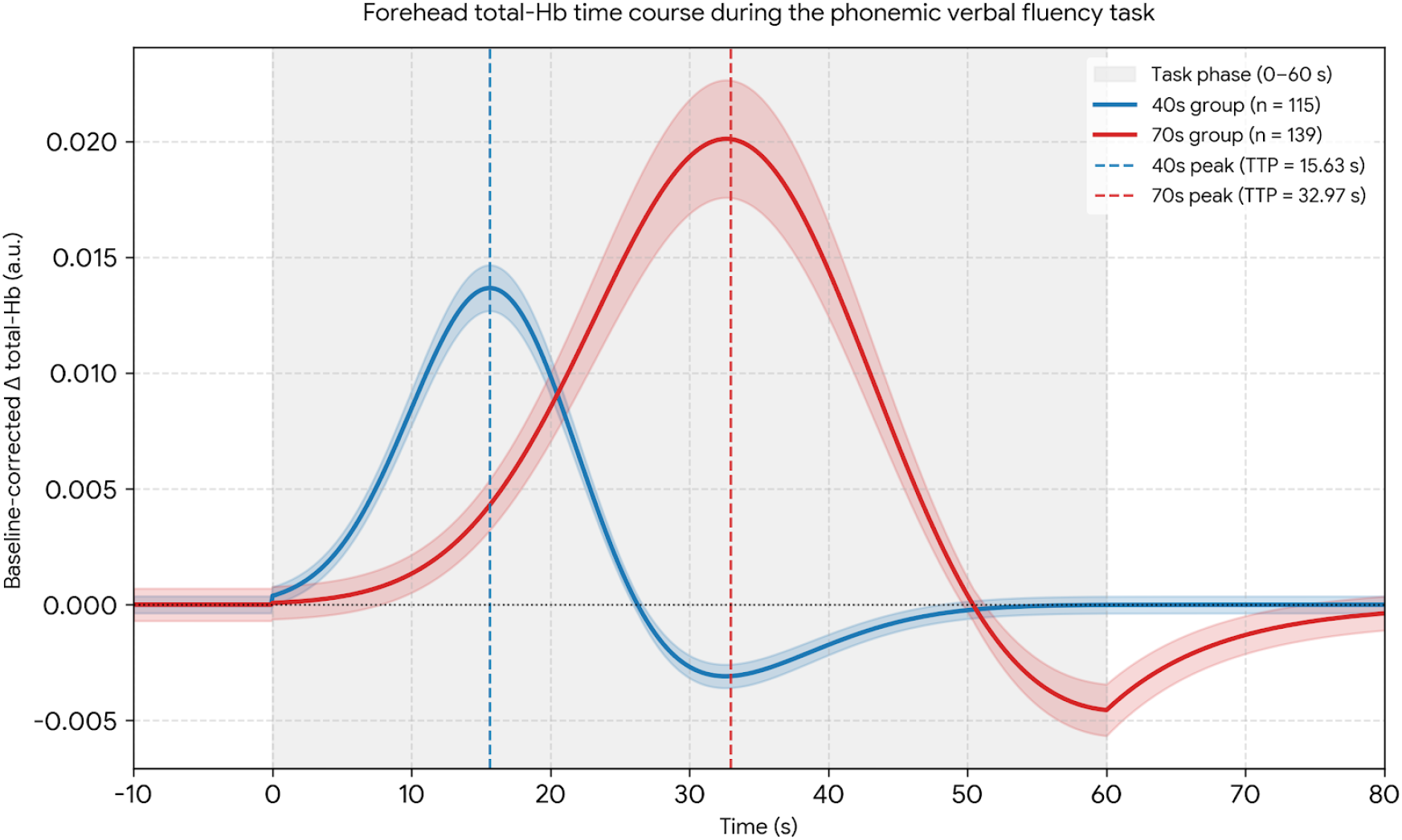
Group-averaged forehead total-hemoglobin (total-Hb) time course. Baseline-corrected total-Hb change for the 40s group (n = 115) and the 70s group (n = 139), recorded with a two-channel, single-wavelength (810 nm) wearable near-infrared spectroscopy device sampling at 10 Hz. Signals were baseline-corrected by subtracting the mean of the 10-s pre-task period (-10 to 0 s) and filtered with a fourth-order zero-phase Butterworth low-pass filter (cutoff 0.1 Hz, forward-backward filtering). The shaded region marks the 60-s task phase (0 to 60 s); the 10-s baseline (-10 to 0 s) and the 20-s recovery period (60 to 80 s) are shown on either side. Solid lines are group means and shaded bands are 95% confidence intervals (mean ± 1.96 × SE). Vertical dashed lines mark the group mean time-to-peak (TTP), 15.63 s for the 40s group and 32.97 s for the 70s group. The vertical axis is relative device output in arbitrary units; values do not represent absolute hemoglobin concentration or localized cortical activity.

Pooled across all participants, TTP was negatively correlated with total word count (r = -0.43, p < 0.001). Age-stratified analyses, however, showed positive correlations in both groups (40s, r = 0.33, p < 0.001; 70s, r = 0.40, p < 0.001), indicating that within each age group participants who produced more words had longer TTP. The positive association persisted after adjustment for age group and sex (partial r = 0.35, p < 0.001), and the age group × TTP interaction was not significant (β = -0.037, 95% CI -0.085 to 0.012, p = 0.137). The reversal in direction between the pooled and within-group correlations is therefore described as an aggregation artifact arising from the large between-group location difference, and we did not conclude that the slope of the TTP-word count association differed by age group. AUC showed no clear within-group association with total word count (40s, r = -0.07, p = 0.481; 70s, r = 0.03, p = 0.710). Scatterplots by age group are shown in Figure 3, and the distribution of AUC is shown in Supplementary Figure S1.

**Figure 3.**
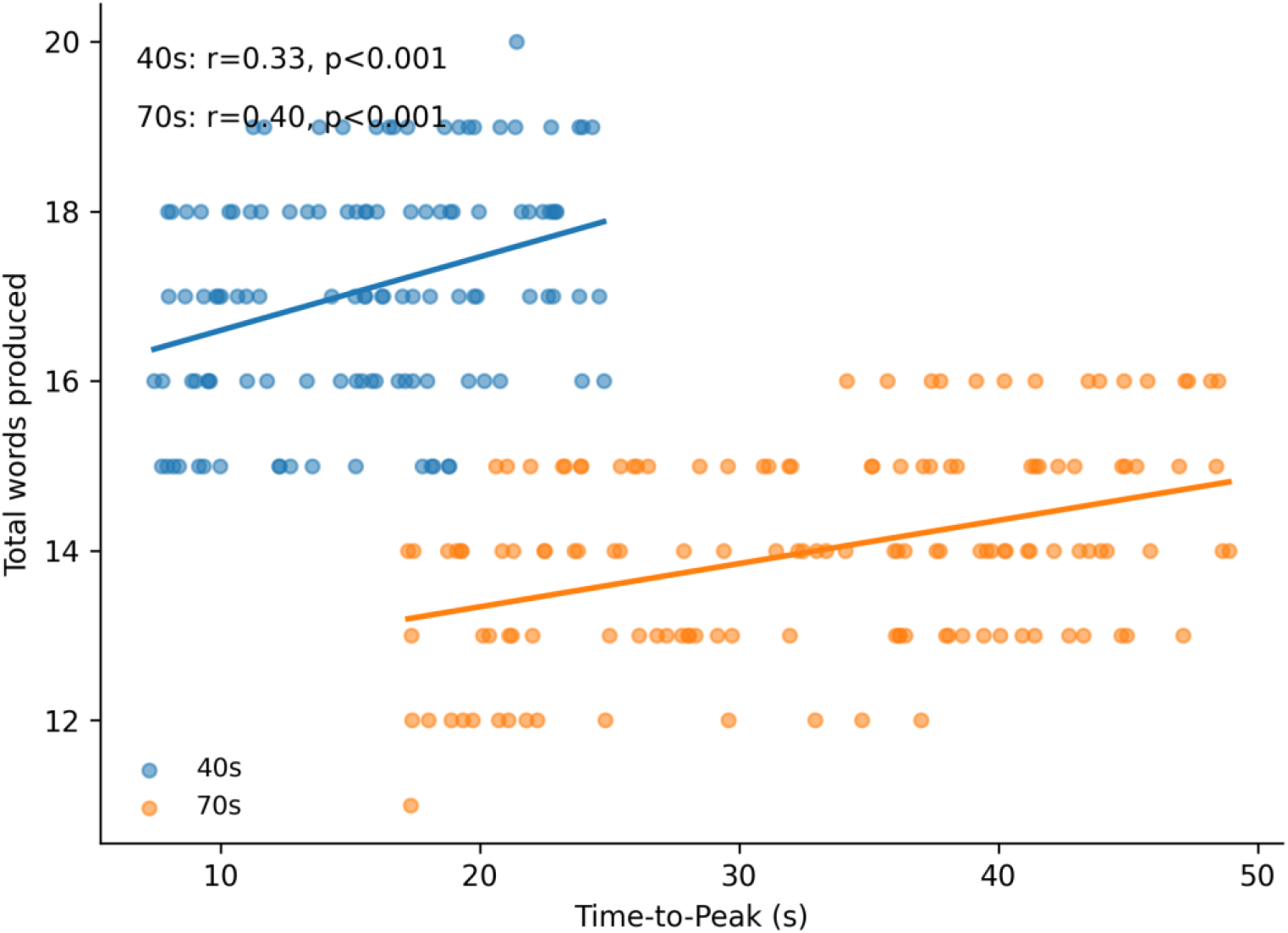
Association between time-to-peak (TTP) and total word count, stratified by age group. Each point represents one participant (40s group, n = 115; 70s group, n = 139). The horizontal axis gives the TTP of the baseline-corrected forehead total-Hb signal in seconds, and the vertical axis gives the total number of valid words produced during the 60-s task. Lines show group-specific ordinary least-squares fits. Within each age group the association was positive (40s, r = 0.33, p < 0.001; 70s, r = 0.40, p < 0.001); it remained positive after adjustment for age group and sex (partial r = 0.35, p < 0.001) and after adjustment for speech offset time (40s, partial r = 0.26, p = 0.006; 70s, partial r = 0.36, p < 0.001). When the two groups were pooled, the correlation reversed in sign (r = - 0.43, p < 0.001). This reversal reflects the large between-group difference in the location of both variables rather than a difference in slope, because the age group × TTP interaction was not significant (β = -0.037, 95% CI -0.085 to 0.012, p = 0.137).

### 3.4 Effect of Speech Duration on total-Hb Measures

Speech offset time was 53.22 ± 5.22 s in the 40s group and 57.27 ± 4.63 s in the 70s group, 4.05 s longer in the 70s group (95% CI 2.82 to 5.28; Hedges’ g = 0.82; p < 0.001). The final production bin was the fourth for 3 participants, the fifth for 72, and the sixth for 40 in the 40s group; the corresponding counts in the 70s group were 1, 36, and 102.

After adjustment for speech offset time and sex, the age-group difference in TTP remained 16.16 s (95% CI 14.13 to 18.19; p < 0.001), only slightly attenuated from the unadjusted 17.34 s. In the mediation analysis, the indirect effect through speech offset time was 1.16 s (bootstrap 95% CI 0.43 to 2.03), accounting for 6.7% of the total effect. For AUC, the adjusted age-group difference was 0.284 a.u.·s (95% CI 0.234 to 0.335; p < 0.001) and the proportion mediated was 0.5%. A subsample of 142 participants continued speaking through the final 10-s bin, so that speech offset time was 60 s in both groups (40s, n = 40; 70s, n = 102). In this subsample, TTP was 16.86 ± 4.71 s and 33.97 ± 9.28 s (Cohen’s d = 2.07, p < 0.001) and AUC was 0.782 ± 0.187 a.u.·s and 1.057 ± 0.203 a.u.·s (d = 1.38, p < 0.001). Both differences were nearly identical to those in the full sample.

Within-group correlations between speech offset time and TTP were weak (40s, r = 0.21; 70s, r = 0.19). TTP occurred on average 37.6 s before speech offset in the 40s group and 24.3 s before speech offset in the 70s group, and no participant showed a peak after speech offset. The positive within-group associations between TTP and total word count also persisted after adjustment for speech offset time (40s, partial r = 0.26, p = 0.006; 70s, partial r = 0.36, p < 0.001).

In summary, relative to the 40s group, the 70s group showed lower verbal fluency performance and longer inter-word pauses but higher AUC and longer TTP in the forehead total-Hb signal. These findings are not direct evidence of localized cortical activity; they indicate that speech behavior and forehead hemodynamic signals manifest differently across age groups.

## 4 Discussion

The principal finding of this study is that, compared with adults in their 40s, adults in their 70s produced fewer words and showed longer inter-word pauses during a phonemic verbal fluency task, while showing higher AUC and longer TTP in the forehead total-Hb signal. Age-group differences in word production and in forehead hemodynamic signals therefore did not run in the same direction. The age-related tendency toward lower fluency performance is consistent with previous reports (Vaughan et al., 2016), although the cross-sectional design means that these differences cannot be read as within-person change. Among the acoustic measures, CPPS was lower in the 70s group, jitter did not differ, and shimmer was also lower in the 70s group. Because acoustic features are affected differently by age, sex, speech task, and analysis conditions depending on the metric (Taylor et al., 2020; Yanushevskaya and Kenny, 2025), these results should not be read collectively as unidirectional deterioration of voice quality.

### 4.1 Temporal Structure of Word Production

The finding that word-production profiles across 10-s bins differed by age group is consistent with reports that within-task variability in verbal fluency carries information distinct from the total word count (Holtzer et al., 2020) and with formal models of age-related change in the efficiency of controlled memory search (Hills et al., 2013). It has also been reported that strategy measures such as clustering and switching do not necessarily add information beyond the total count in phonemic tasks (Haugrud et al., 2011). The bin-specific counts and mean inter-word pause durations used here are therefore best regarded as direct measures of the temporal structure of speech behavior rather than as equivalents of conventional clustering and switching indices. Notably, word production in the final 10-s bin fell to 0.70 words in the 40s group but remained at 1.48 words in the 70s group, indicating a steeper late-task decline in the younger group. This difference in temporal distribution is informative when interpreting the age-group difference in TTP, although bin-specific counts alone cannot establish how long lexical search continued.

### 4.2 Forehead total-Hb Responses

With respect to TTP, the 70s group showed values that were on average 17.34 s longer, while within each age group producing more words was positively associated with longer TTP. The age group × TTP interaction was not significant, so there was no evidence that the slope of this association differed by age group. BOLD-fMRI studies report prolongation of the time-to-peak of the hemodynamic response function with healthy aging (West et al., 2019), but that measure is not identical to a TTP derived from single-wavelength total-Hb. Associations between hemodynamic latency and cognitive function (Anderson et al., 2020) and age-related changes in neurovascular coupling (Mukli et al., 2024) have also been reported. TTP in the present study should therefore not be equated with slower neural processing; it is better interpreted as a measure that may reflect both the temporal structure of task performance and age-dependent hemodynamic characteristics.

Because forehead total-Hb signals can be influenced by speech-related respiratory fluctuation, scalp blood flow, and systemic hemodynamics, it was necessary to test whether the age-group differences in TTP and AUC simply reflected differences in how long speech continued. The between-group difference in speech offset time was only 4.05 s, and adjusting for it changed the TTP difference from 17.34 s to 16.16 s. The proportions mediated by speech offset time were 6.7% for TTP and 0.5% for AUC, and the between-group differences were nearly unchanged in the subsample matched on speech offset time. TTP also occurred well before speech offset, and within-group correlations between the two were weak. The observed age-group differences in forehead total-Hb responses were therefore not explained by the duration of continued speech. This verification nevertheless used an upper-bound approximation of speech offset based on 10-s resolution, and confirmation with finer-grained speech timing is needed.

Although the 70s group produced fewer words, it showed a higher AUC of the forehead total-Hb signal. The HOT-2000 is a single-wavelength device, however, and the total-Hb signal may include scalp blood flow and systemic hemodynamic contributions in addition to cortical neurovascular responses. No clear within-group association was found between AUC and total word count. The higher AUC in the older group therefore cannot be equated with better cognitive performance, greater neural activity, or a compensatory mechanism, and is better read as indicating that the response pattern of forehead hemodynamic signals during task performance differs across age groups.

### 4.3 Dissociation Among Measures and Implications for Characterizing Aging

MoCA-J total scores clustered near the ceiling in both groups yet showed a small statistically significant age-group difference, whereas the continuous measures derived from the same task differed more substantially and in different patterns. The contribution of this study therefore lies not in demonstrating change that the MoCA-J “failed to detect,” but in describing age-group differences in speech behavior and hemodynamic response on scales distinct from the total score. Because the phonemic verbal fluency task analyzed here is a component of the MoCA-J, it is not independent of the total score, and these findings cannot be generalized to claims about improved screening accuracy or early detection of dementia.

The pattern of results is nevertheless informative for how aging in speech might be characterized. The measures obtained from one 60-s speech sample did not move together. Word count, inter-word pause duration, and the within-task production gradient - measures plausibly weighted toward central, cognitive-linguistic contributions - differed markedly between groups. Among the acoustic measures, which are weighted toward peripheral, phonatory and respiratory contributions, CPPS differed between groups whereas jitter did not, and shimmer differed in the direction opposite to that expected from aging. TTP and AUC, both derived from the same total-Hb signal, also behaved differently: TTP was associated with word count within each age group whereas AUC was not. These dissociations are consistent with the view that different biological and functional systems change at different rates and in different directions within individuals (Ahadi et al., 2020) and that a single summary index, whether a screening score or a single voice metric, is unlikely to capture aging in speech adequately.

Two implications follow. First, characterizing aging in speech will require the joint modeling of several measurement axes rather than the refinement of any one metric. Concurrent measurement of physical function, such as grip strength, gait speed, and frailty status, would allow the peripheral contribution to be estimated directly rather than inferred, and would test whether acoustic and temporal measures load on separable axes. Second, because the present design is cross-sectional and compares only two age bands, the observed contrasts cannot distinguish between-person differences from within-person change (Salthouse, 2019). Longitudinal follow-up of the same individuals with repeated speech, cognitive, and physical-function measurement is required to estimate individual trajectories along each axis and to determine whether the axes diverge within persons as they do between groups here (Elliott et al., 2026).

### 4.4 Limitations

This study has several limitations. First, it was a cross-sectional comparison between adults in their 40s and 70s, so the observed differences cannot be interpreted as within-person change, and a continuous age gradient cannot be estimated from two age bands (Salthouse, 2019).

Second, the 40s group comprised care-house staff and the 70s group comprised care-house residents, so age group could not be fully separated from participant role and living environment. Although procedures and environment were standardized, residual confounding from this recruitment structure cannot be excluded. Participants in both groups self-selected under recruitment criteria restricted to healthy individuals, and the 70s group was further restricted by care-house admission requirements to older adults independent in activities of daily living. Observed ranges of total word count were correspondingly narrow (15-20 words in the 40s group, 11-16 words in the 70s group), with standard deviations of 1.33 and 1.18 words, clearly smaller than those reported in normative verbal fluency studies of community populations. This sample was therefore strongly selected toward better health, and variability in cognitive function and fluency performance was almost certainly smaller than in the general population. The effect sizes reported here are valid only for similarly selected samples and should not be extrapolated to the general older population.

Third, MoCA-J scores were strongly concentrated at the ceiling, with perfect-score rates of 95.7% and 82.7%, markedly higher than previously reported for healthy older adults. This most likely reflects the selection described above, although an influence of administration or scoring procedure cannot be excluded. The MoCA-J was administered by a single researcher using a standardized procedure, but because age group was inherently apparent, blinding of the examiner and scorer was not possible and no independent rescoring was performed. Although the age-group difference was significant by both tests, the median and interquartile range were 30 [30-30] in both groups, so the practical magnitude of the difference should be interpreted cautiously.

Fourth, the phonemic verbal fluency task analyzed here contributes to the MoCA-J total score and is therefore not independent of it, although the sensitivity analysis using the modified score excluding that item showed no age-group difference at the two-sided 5% level. Fifth, years of education were not collected, so residual confounding from educational history or lexical-use experience cannot be excluded, and the education-related point adjustment in the MoCA scoring rules could not be applied. Sixth, although individuals requiring regular treatment for hypertension, diabetes, cardiovascular disease, or comparable conditions, and those using vasoactive medication, were excluded at recruitment, blood pressure and vascular physiological measures were not collected as continuous variables, so vascular contributions could not be fully evaluated. Seventh, the task consisted of a single trial with the single syllable /ka/, and reproducibility across other cue sounds or repeated trials was not examined.

Eighth, the HOT-2000 is a two-channel, single-wavelength device that cannot separate oxygenated from deoxygenated hemoglobin. It lacks short-separation channels, and quantitative artifact correction using accelerometry or pulse signals was not performed, so the total-Hb signal may contain systemic and extracerebral components including scalp blood flow, cardiac activity, and respiration (Sato et al., 2013; Yücel et al., 2021). Although no abrupt head movement or pulse increase was observed during measurement, these influences were not quantitatively removed. AUC and TTP should accordingly be interpreted as task-related forehead total-Hb responses rather than localized cortical activity.

Ninth, speech was recorded with a microphone approximately 2 m from the participant across a desk. This distance departs substantially from the close-microphone placement recommended for voice assessment, typically 5 to 15 cm from the mouth, and is vulnerable to reduced signal-to-noise ratio and room reverberation. CPPS, jitter, and shimmer all depend on recording distance, signal-to-noise ratio, and vocal sound-pressure level, so the absolute values reported here cannot be compared directly with normative values from other studies. The jitter (approximately 2.2%) and shimmer (approximately 20%) values obtained were substantially higher than levels commonly reported for healthy adults, indicating that period extraction in connected speech was not stable, and the finding that shimmer was lower in the 70s group runs counter to the direction expected from aging and may be an artifact of the recording conditions. Jitter and shimmer should therefore be treated as reference values only. For CPPS, a systematic influence mediated through signal-to-noise ratio cannot be excluded if vocal sound-pressure levels were lower in the 70s group; because sound-pressure level and signal-to-noise ratio were not quantified, this possibility could not be evaluated.

Tenth, speech offset time was an upper-bound approximation based on 10-s resolution and therefore falls later than the actual end of the final word; the stability of the results under finer temporal quantification has not been established. Because the latency to the first word and the durations of individual words were not measured directly, the time budget of the 60-s task could not be fully decomposed.

## 5 Conclusion

Between adults in their 40s and 70s we observed distinct patterns of group difference in the amount and temporal structure of word production during a phonemic verbal fluency task, in acoustic features, and in forehead total-Hb responses. The 70s group produced fewer words and showed longer inter-word pauses, while showing longer TTP and higher AUC; these hemodynamic differences were not explained by how long speech continued during the task. Critically, the measures did not move together: temporal measures of word production, acoustic measures of voice quality, and the magnitude and timing of the hemodynamic response dissociated from one another and from the screening total score. Even in a group whose MoCA-J scores clustered at the ceiling, a single 60-s speech sample thus yielded several partially independent descriptions of age-group difference. Because the sample was selectively recruited from healthy individuals and single-wavelength total-Hb signals do not index localized neural activity, these results are descriptive rather than mechanistic. They indicate, however, that characterizing aging in speech will require longitudinal, individual-level modeling along multiple measurement axes, with concurrent assessment of physical function so that peripheral and central contributions can be estimated separately rather than inferred from a single index.

## 7 Author Contributions

KN: Conceptualization, Methodology, Investigation, Data curation, Formal analysis, Visualization, Writing - original draft, Writing - review and editing, Funding acquisition, Project administration, Supervision.

## 9 Acknowledgments

The author thanks Keiichi Yamazaki (Saitama University) and Akiko Yamazaki (Tokyo University of Technology) for their academic advice, discussion, and support throughout this study.

## 10 Ethics Statement

The studies involving humans were approved by the Ethics Committee of Gunma University Graduate School of Medicine (approval no. HS2024-273; approved April 2024). The studies were conducted in accordance with the local legislation and institutional requirements. The participants provided their written informed consent to participate in this study. Written informed consent was obtained for the publication of any potentially identifiable images or data included in this article.

## 11 Generative AI Statement

We do not use generative AI.

## 12 Supplementary Material

Supplementary Figure S1 shows the distribution of AUC of the forehead total-Hb response by age group. Supplementary Figure S2 shows the participant recruitment and analysis flow. Both are provided as separate files.

## 13 Data Availability Statement

The raw datasets are not publicly available because they contain potentially identifiable voice recordings. The de-identified derived dataset supporting the analyses reported here (participant-level word counts, bin-specific counts, pause and speech-offset measures, acoustic summaries, and NIRS summary measures) is available from the corresponding author on reasonable request, subject to approval by the relevant ethics committee and institutional requirements.

---

**Supplementary Figure S1.**
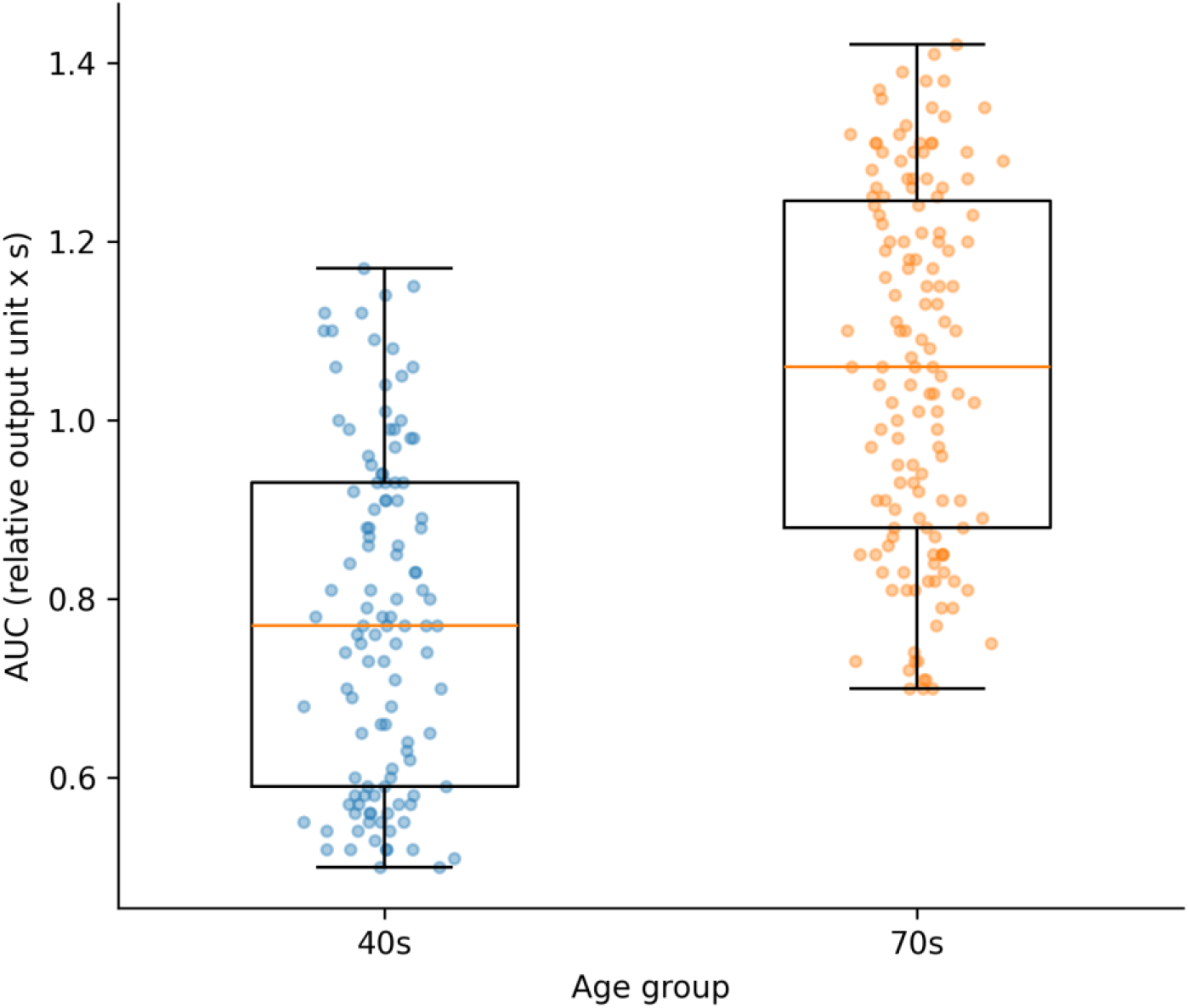
Distribution of the area under the curve (AUC) of the forehead total-Hb response by age group. AUC was obtained by integrating the baseline-corrected total-Hb change from task onset to 60 s and is expressed in arbitrary units multiplied by seconds (a.u.·s). Values were 0.776 ± 0.188 a.u.·s in the 40s group (n = 115) and 1.059 ± 0.203 a.u.·s in the 70s group (n = 139), a difference of 0.283 a.u.·s (95% CI 0.235 to 0.332; Hedges’ g = 1.44; FDR-adjusted q < 0.001). Because the device reports relative change, AUC is an integrated device-output measure and is interpreted as an exploratory hemodynamic summary rather than an absolute measure of cerebral blood flow. AUC showed no clear within-group association with total word count (40s, r = -0.07, p = 0.481; 70s, r = 0.03, p = 0.710).

**Supplementary Figure S2.**
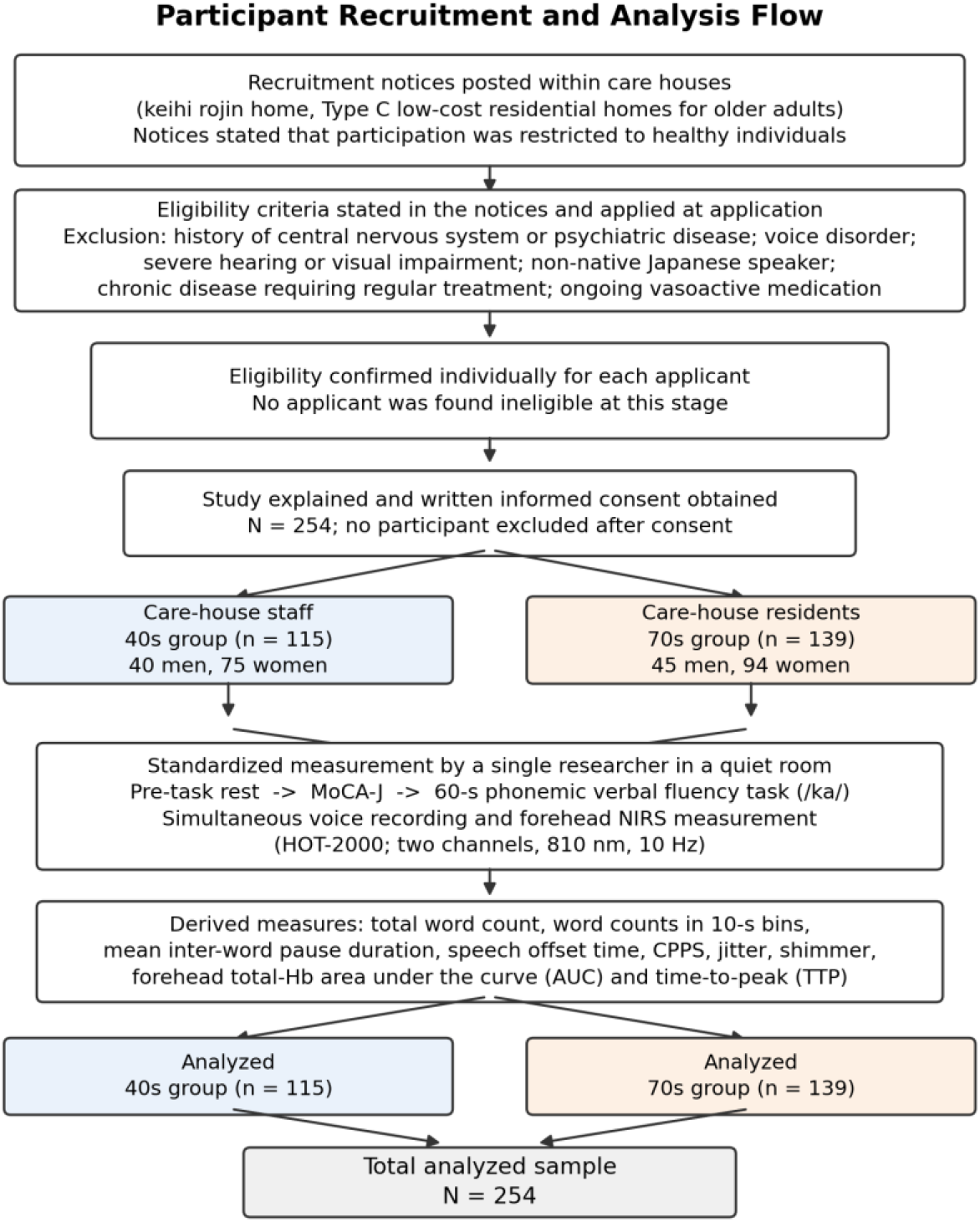
Participant recruitment and analysis flow. Participants were recruited through notices posted within care houses (keihi rojin home, Type C low-cost residential homes for older adults in Japan). The 40s group comprised care-house staff and the 70s group comprised residents of care houses of the same type. Eligibility criteria restricting participation to healthy individuals were stated in the recruitment notices and confirmed individually for each applicant before the study was explained. No applicant was found ineligible at the eligibility-confirmation stage and no participant was excluded after consent, so all 254 individuals who provided written informed consent were analyzed (40s group, n = 115; 70s group, n = 139). All participants underwent the same standardized protocol in a quiet room after a pre-task rest period, administered by a single researcher.

